# Integrating multi-network topology for gene function prediction using deep neural networks

**DOI:** 10.1101/532408

**Authors:** Hansheng Xue, Jiajie Peng, Xuequn Shang

## Abstract

**Motivation:** The emerging of abundant biological networks, which benefit from the development of advanced high-throughput techniques, contribute to describing and modeling complex internal interactions among biological entities such as genes and proteins. Multiple networks provide rich information for inferring the function of genes or proteins. To extract functional patterns of genes based on multiple heterogeneous networks, network embedding-based methods, aiming to capture non-linear and low-dimensional feature representation based on network biology, have recently achieved remarkable performance in gene function prediction. However, existing methods mainly do not consider the shared information among different networks during the feature learning process. Thus, we propose a novel multi-networks embedding-based function prediction method based on semi-supervised autoencoder and feature convolution neural network, named *DeepMNE-CNN*, which captures complex topological structures of multi-networks and takes the correlation among multi-networks into account.

**Results:** We design a novel semi-supervised autoencoder method to integrate multiple networks and generate a low-dimensional feature representation. Then we utilize a convolutional neural network based on the integrated feature embedding to annotate unlabeled gene functions. We test our method on both yeast and human dataset and compare with four state-of-the-art methods. The results demonstrate the superior performance of our method over four state-of-the-art algorithms. From the future explorations, we find that semi-supervised autoencoder based multi-networks integration method and CNN-based feature learning methods both contribute to the task of function prediction.

**Availability:** *DeepMNE-CNN* is freely available at https://github.com/xuehansheng/DeepMNE-CNN

## 1 Introduction

With the rapid development of high-throughput experimental techniques, the quality and variety of biological data have experienced an exponential increase during the past decades. This phenomenon has posed new challenges for biologists to effectively extract and comprehensively understand intrinsic relation information cross various data sources. Thus, many approaches have been proposed to integrate multiple data sources to improve the effectiveness on many tasks (Zitnik *et al.*, 2018a), such as protein function prediction (Cozzetto *et al.*, 2013; Wass *et al.*, 2012), drug-target interaction prediction (Zitnik *et al.*, 2018b) and gene function prediction (Re and Valentini, 2010).

Accurate annotation of gene function is one of the most important and challenging problems in biological area. Annotating gene function, also termed as gene function prediction, aims to assign an unknown gene to the correct functional category in the annotation database, such as Gene Ontology. To solve this problem, lots of methods based on different types of biological data have been proposed, such as amino acid sequence-based method (Clark and Radivojac, 2011), protein structure-based method (Pal and Eisenberg, 2005) and gene expression-based method (Huttenhower *et al.*, 2006). With the improvement of experimental methods, several types of association between genes could be obtained, such as gene co-expression, genetic interaction. The complex associations between genes are usually represented in terms of gene association networks, such as gene co-expression network Stuart *et al.* (2003) and genetic interaction network Baryshnikova *et al.* (2013). Generally, each type of network represents a kind of association between genes. Based on gene association networks, several gene or protein function prediction methods have been proposed (Lehtinen *et al.*, 2015; Roded *et al.*, 2007). Recently, network-based gene function prediction has been ushered into a new era by integrating multiple networks. Specifically, multi-network-based function prediction has been proved better than single-network-based methods (Re and Valentini, 2010; Cozzetto *et al.*, 2013; Wass *et al.*, 2012), because of the complementary nature of different data sources. Thus, several algorithms have been proposed for gene function prediction by integrating multiple biological networks (Cho *et al.*, 2016; Mostafavi *et al.*, 2008; Cao *et al.*, 2014).

Most of existing multi-networks based gene function prediction methods mainly contain two components: integrating multiple functional association networks into a single network and utilizing classification algorithms (such as support vector machine (Chang and Lin, 2011)) to label unannotated genes based on the ensemble network.

Several approaches focus on generating an integrated network by combing information from multiple networks, such as probabilistic methods, like Bayesian inference (Franceschini *et al.*, 2013; Lee *et al.*, 2011; Wong *et al.*, 2015), kernel-based methods (Yu *et al.*, 2015) or weighted averaging or summing (Mostafavi *et al.*, 2008; Lanckriet *et al.*, 2004; Tsuda *et al.*, 2005). For instance, a weighted-sum method fuses networks by assigning weights for different individual networks. The weight of each network relies on its predictiveness of a set of positively labeled genes that have the same specific function (Lanckriet *et al.*, 2004; Tsuda *et al.*, 2005). However, these methods prone to overfitting because some functional categories have only a few annotations (Sara and Quaid, 2010). GeneMANIA (Sara and Quaid, 2010; Mostafavi *et al.*, 2008) is a network-based gene function prediction method which solves a constrained linear regression problem by integrating various networks into a single kernel, and this method utilizes Gaussian label propagation on the resulting kernel to annotate unlabeled genes. Similarity Network Fusion (SNF) (Wang *et al.*, 2014) is a widely used networks integration method, which constructs gene association network for each available data source and then efficiently fuses these networks into one integrated network. The core of SNF is merging multi-networks into a single network by taking advantages of the complementarity of all networks. However, these methods may lead to information loss problem in the process of summarizing multiple networks into a single one (Cho *et al.*, 2016). In contrast, some methods try to train individual classifiers on different networks and combine these predictions to a final result using ensemble learning methods (Yan *et al.*, 2010; Yu *et al.*, 2013; Hansen *et al.*, 2018; Valentini, 2014). However, these approaches do not consider the correlations among different networks during the model training process. In addition, such methods often suffer from time and machine equipment constraints.

Mashup (Cho *et al.*, 2016) is an integrative and scalable framework for capturing low-dimensional feature representations of genes from multiple networks constructed from various data sources. It utilizes a matrix factorization-based approach on a collection of gene interaction networks to obtain compact and low-dimensional feature vectors of genes. These feature vectors can describe the internal and hidden features across all networks for genes. Then, Mashup trains a support vector machine (SVM) classifier to calculate functional probabilistic distribution for unannotated genes based on the obtained low-dimensional feature representations. The key of Mashup is the feature learning from multi-networks. Node feature extraction based on multi-networks has been proved useful and effective on many tasks, including gene function prediction. However, matrix factorization-based method (e.g. singular value decomposition) used in Mashup is a linear and shallow approach which would be difficult to capture complex and highly non-linear structure across all networks.

In addition, Diffusion State Distance (Cao *et al.*, 2014) is a diffusion-based method that uses random walk with restart to capture the local topology for each gene. The idea is that genes with similar diffusion states should have similar function. Collective Matrix Factorization (Zitnik *et al.*, 2015) is a kind of matrix factorization-based approach, which jointly factorizes multiple networks to obtain a latent representation of genes. Similar to Mashup, these feature representations could be used in the following classification mode. However, these approaches both cause information loss when summarizing multiple networks into a single one.

Due to information loss within multi-networks integration and complex non-linear features across multi-networks, it is significant but challenging to generate feature representations of genes based on multi-network integration. Learning network topology features and generating low-dimensional vector representations can be described as network representation learning or network embedding. Most existing network embedding methods aims to learn feature representation from single network, such as node2vec (Grover and Leskovec, 2016), DeepWalk (Perozzi *et al.*, 2014) and LINE (Zhang *et al.*, 2015). However, to the best of our knowledge, no existing method is designed for learning feature representation of genes from multiple networks, although there is urgent requirement in related area.

During past decades, Deep Learning has been widely used in image processing (Krizhevsky *et al.*, 2012; Karpathy *et al.*, 2014), natural language processing (Kim, 2014) and achieved remarkable results. Recently, researchers have tried to apply deep learning models on the areas of bioinformatics (S *et al.*, 2016), such as gene ontology annotation (Chicco *et al.*, 2014), miRNA-disease association (Peng *et al.*, 2018), regulatory genomics and cellular imaging analysis (Angermueller *et al.*, 2016). Deep learning can solve the high dimensional feature learning problems effectively by the non-linear activation function. A few approaches have been proposed recently to learn non-linear network representations from complex data sources using autoencoder and convolutional neural network (Tian *et al.*, 2014; Cao *et al.*, 2016; Zitnik *et al.*, 2018b; Gligorijevic *et al.*, 2017). AutoEncoder (Rumelhart *et al.*, 1986; Baldi, 2011) is a typical unsupervised deep learning model, which aims to learn a new encoding representation of input data. Because of its superiority on dimensionality reduction and feature extraction, autoencoder is often used on graph clustering (Tian *et al.*, 2014) and medical image search (Sharma *et al.*, 2016). Convolutional neural network (Krizhevsky *et al.*, 2012) is a typical feed-forward artificial neural network, which has been commonly applied on computer vision and natural language processing (Ronan Collobert, 2008). Besides, CNN is also introduced in drug discovery (Wallach *et al.*, 2015) and achieved promising results.

Although autoencoder and convolutional neural network have been proved useful end effective in learning biological network topology information, none of existing model is designed for multi-network embedding specifically. In this paper, we propose a novel multi-networks embedding approach and use it to annotate unlabeled gene function, named DeepMNE-CNN. DeepMNE-CNN mainly contains two components. One component is multi-networks integration framework, which applies a novel semi-supervised autoencoder to map input networks into a low-dimension and non-linear space based on prior information constraints. The other is CNN-based function predictor, which use convolutional neural network to learn feature embedding. Here are the major contributions:

- To incorporate prior information as constraints, we propose a novel semi-supervised autoencoder model, DeepMNE, to obtain compact and non-linear topological feature representations of genes based on multiple networks.
- We utilize a convolutional neural network (CNN) architecture to predict gene function based on the feature vectors summarized from multiple networks.
- The evaluation results show that DeepMNE-CNN outperforms the existing state-of-the-art methods on the task of gene function prediction, and multi-networks integration framework implies promising priority comparing with the others.

## 2 Methods

DeepMNE-CNN contains two main parts, integrating multi-networks based on revised autoencoder and predicting gene function using CNN. Multi-networks integration can be described as a semi-supervised feature learning algorithm, which considers constraints extracted from other networks in multi-network embedding. Then, we utilize convolutional neural network to predict gene function based on the output of DeepMNE. The workflow of DeepMNE-CNN is shown in Figure 1.

**Fig. 1.**
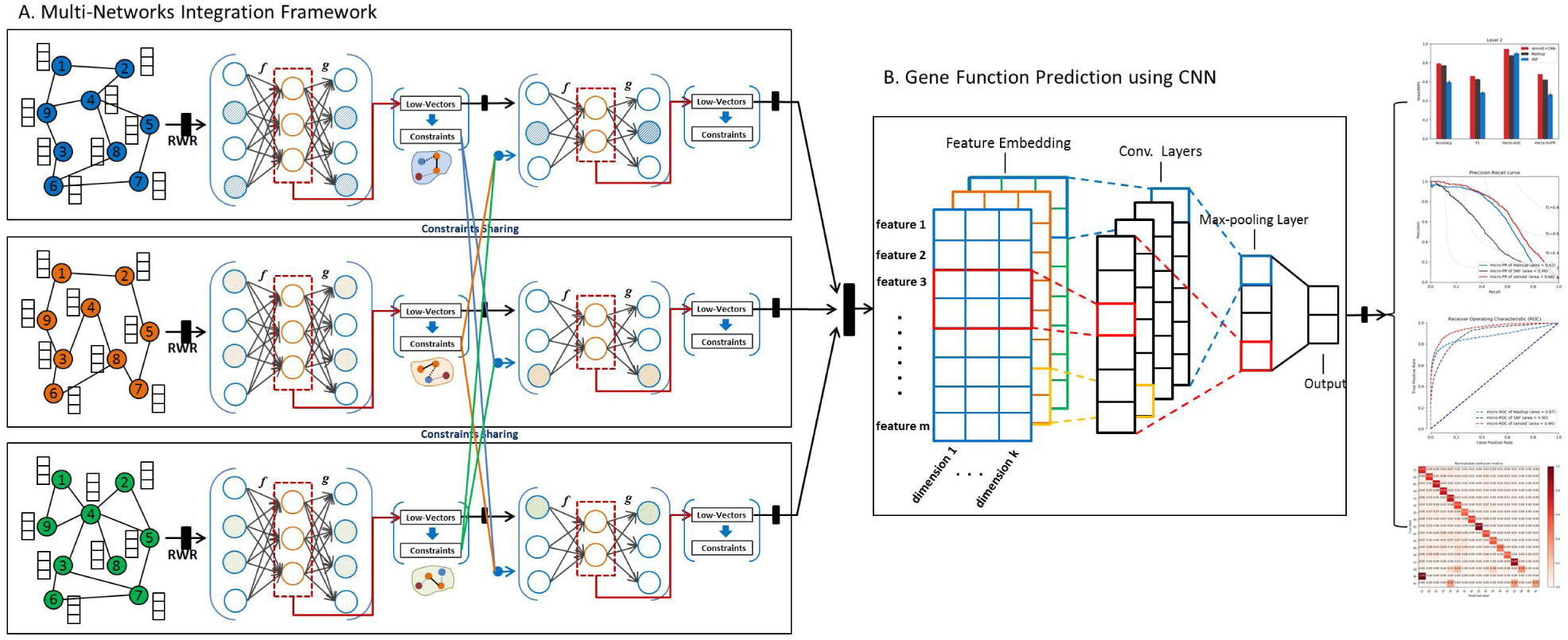
The structure of DeepMNE-CNN algorithm. This framework mainly contains two parts, multi-networks integration framework (DeepMNE) and CNN-based gene function prediction. For DeepMNE, we first run random walk with restart (rwr) to learn global structure of networks. Then, constraints extraction and application with semi-supervised autoencoder are iteratively implemented on DeepMNE framework to integrate multi-networks. After obtaining the integrated representations of multi-networks, we can train convolution neural network based on the outputs of DeepMNE to annotate gene function categories. **Fig. 2.** DeepMNE-CNN improves gene function prediction performance in yeast. We compared the function prediction performance of DeepMNE-CNN to other state-of-the-art approaches, Mashup and SNF in yeast. Figure represents the three different levels of functional categories respectively.

Given *k* networks that include the same set of nodes *V* but different connectivity between nodes, labeled as {*G*^(1)^, *G*^(2)^,*…, G*^(*k*)^}, a specific network *G*_*i*_, each network is represented as *G*^(*i*)^ = (*V, E*^(*i*)^), where *i* ∈ {1, 2, *…, k*}. Let *V* be a set of *n* nodes {*v*_1_, *v*_2_, *…, v*_*n*_} in a multi-networks set. Let *E* be a set if edges between pairs of *n*-nodes {*v*_1_, *v*_2_, *…, v*_*n*_} in a specific single network. Our aim is to learn a low-dimension feature representation for each *v ∈ V* based on the topological information contained in {*G*^(1)^, *G*^(2)^,*…, G*^(*k*)^}.

### 2.1 Multiple networks integration framework

In this section, we propose a novel multi-network embedding algorithm, termed as DeepMNE. The main framework is a DNN structure with autoencoder (AE) and Semi-Supervised autoencoder (semiAE) as its building block. The whole process includes three parts: learning single network topology information, extracting prior constraints and integrating constraints with semiAE model. The main framework of DeepMNE is shown in Figure 1.A.

Specifically, the first layer of DeepMNE framework is the original autoencoder, which is used for feature extraction and dimension reduction. Starting from the second layer, a revised autoencoder (semiAE) is used for constraint integration and dimension reduction. The dimension of input networks decreases constantly with the extension of the whole iteration model.

#### Step 1: Learning the global structure of single network based on RWR

It has been proved that random walk with restart (RWR) could capture global associations between nodes in a network (Cho *et al.*, 2016). Instead of inputting adjacency matrices into DeepMNE directly, we run RWR on each network to capture single network topological information and convert it into feature representations of nodes. The adjacency matrix only describes the relationships between any directly connected nodes, ignoring the global structure of a network. RWR can overcome this drawback, and represent nodes using these high-dimensional network structural information. Besides, we choose the RWR method instead of other recently proposed network embedding approaches, such as node2vec (Grover and Leskovec, 2016) and DeepWalk (Perozzi *et al.*, 2014), to capture the topological information, because these state-of-the-art algorithms are computationally intense and require additional hyper-parameter fitting (Gligorijevic *et al.*, 2017).

Let *M*^*k*^ denote the adjacency matrix of the *k–*layer network *G*^(*k*)^ = (*V, E*^(*k*)^). The RWR from node *v*_*i*_ can be described as the following recurrence relation.

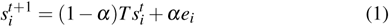

where *α* is the restart probability; *e*_*i*_ is a n-dimensional initial feature vector; 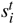 is a n-dimensional feature vector of gene *i*, and each entry indicates the probability of a gene being visited after *t* steps in the random walk; *T* is the transition probability matrix, and each entry *T*_*i* *j*_ saves the probability from gene *j* to gene *v*_*i*_, which can be calculated as 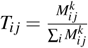 Based on RWR, we can obtain a matrix *S*, in which *S*_*i* *j*_ is the relevance score between node *v*_*i*_ and *v* _*j*_ defined by RWR-based steady state probabilities.

#### Step 2: Extracting prior constraints

The idea of constraints comes from semi-supervised clustering. The pairwise constraints can be typically formatted as must-link and cannot-link constraints (Basu *et al.*, 2004). The pairwise constraints can be described as follows: a must-link constraint indicates that these nodes are highly similar or belong to the same cluster, while a cannot-link constraint indicates that two points in the pair are highly dissimilar or belong to different clusters.

In our model, we calculate pearson correlation coefficient (PCC) value to measure the pairwise similarity between gene nodes. Given two gene feature vectors *A* and *B*, the pcc value of these two genes can be calculated as:

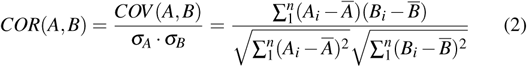

After calculating pcc value of all pairs of genes, we use two strategies to extract constraints. One is to calculate and sort pairwise pearson correlation coefficient (PCC) of all pairs of nodes based on their feature vectors. The top-*k* pairs are considered as the must-link constraints and the bottom-*k* pairs are considered as the cannot-link constraints. The other is to set two thresholds for must-link and cannot-link, labeled as *f*_1_ and *f*_2_ respectively. In detail, a pairs can be adopted as a must-link constraint if its PCC value is larger than *f*_1_, and a pair is considered as cannot-link constraint if the PCC value is smaller than *f*_2_.

After extracting the constraints from the previous layer (*i* layer), we can apply the must-link and cannot-link constraints to the next layer (*i*+1 layer) as the prior information.

#### Step 3: Integrating constraints using Semi-supervised AutoEncoder

The key question of DeepMNE is how to integrate prior constraints into the network representation through autoencoder. We revise the original autoencoder and propose a novel variant of autoencoder, termed as Semi-Supervised AutoEncoder (semiAE). Starting from the second layer, the input includes both low-dimensional representations and constrains from previous layer. It is noted that constraints from previous layers’ building blocks are based on different networks. Therefore, constraints may be conflicting. To solve this problem, we would merge these constraints and take the intersection of all the constraints as the input of semiAE.

Autoencoder is an unsupervised model which is composed of two parts, i.e. the encoder and decoder. The encoder operation transforms the input high-dimensional data into a low-dimensional feature representation, and a similar “decoder” operation to recover the input data from the low-dimensional feature vectors. The low-dimensional code is then used as a compressed representation of the original data. Let *x*_*i*_ be the *i*- th input vector or node representation of network, and *f* and *g* be the activations of the hidden layer and the output layer respectively. We have *h*_*i*_ = *f* (*Wx*_*i*_ + *b*) and *y*_*i*_ = *g*(*Mh*_*i*_ + *d*), where Θ = {*θ*_1_, *θ*_2_} = {*W, b, M, d*} are the parameters to be learned, *f* and *g* are the non-linear operators such as the sigmoid function (*sigmoid*(*z*) = 1*/*(1 + *exp*(*–z*))) or tanh function (*tanh*(*z*) = (*e*^*z*^ – *e*^*−z*^)*/*(*e*^*z*^ + *e*^*−z*^)). Then the optimization goal is to minimize the reconstruction error between the original data *x*_*i*_ and the reconstructed data *y*_*i*_ from the new representation *h*_*i*_.

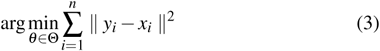

The original autoencoder cannot model the constraints obtained from previous layers. We propose semiAE to take these constraints into account. Let *M* be a set of must-link pairwise constraints where (*x*_*i*_, *x* _*j*_) *∈ M* implies the strong association between *x*_*i*_ and *x* _*j*_. Let *C* be a set of cannot-link pairwise constraints where (*x*_*i*_, *x* _*j*_) *∈ C* implies *x*_*i*_ and *x* _*j*_ are unrelated. The number of constraints is much less than the size of the network *|M|* + *|C| ≤ |S|*.

The hypothesis is that *x*_*i*_ and *x* _*j*_ should also close based on the low-dimensional space if there is a must-link constraint between them in previous layer. Ideally, after encoding, two must-link nodes should be closer, and two cannot-link nodes may be more distant. Mathematically, let *d*(*h*(*x*_*i*_), *h*(*x*_*j*_)) be the error score (difference) between *x*_*i*_ and *x* _*j*_ in the encoded space. For Must-link, *d*(*x*_*i*_, *x* _*j*_) should be larger than *d*(*h*(*x*_*i*_), *h*(*x*_*j*_)); for Cannot-link, *d*(*x*_*i*_, *x* _*j*_) should be smaller than *d*(*h*(*x*_*i*_), *h*(*x*_*j*_)). If the pair (*x*_*i*_, *x* _*j*_) is a must-link constraint, we add a penalty on the loss function. Similarity, if the pair (*x*_*i*_, *x* _*j*_) is a cannot-link constraint, we add a reward on the loss function. The loss function for modeling constraints is defined as follows:

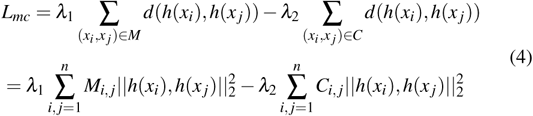

where matrix *M* and *C* are extracted must-link and cannot-link constraint matrix from previous layer respectively; *h*(*x*_*i*_) and *h*(*x*_*j*_) are hidden layer representation of input gene representations *x*_*i*_ and *x* _*j*_; *λ*_1_ and *λ*_2_ are weight coefficient, controlling the influence of penalty and reward respectively.

To combine constraints with autoencoder, we propose a novel semi-supervised autoencoder, which integrates Eq. (3) and Eq. (4) and joint minimizes the following objective function:

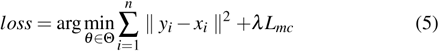

The first part of Equation 5 measures the squared error between input and output node features, and the second part measures error score of constraints in hidden layer.

### 2.2 Gene Function Prediction with CNN

After obtaining low-dimensional feature representations by DeepMNE, we train a convolutional neural network (CNN) model to annotate gene function. The whole structure of CNN-based gene function prediction is shown in Figure 1.B.

Let *E* ∈ ℝ ^*n×m×k*^ be the low-dimensional network representation, which obtain from multi-networks integration framework, DeepMNE. *n, m* and *k* represent the number of genes, the length of gene feature embedding, and the number of networks respectively. Let *x*_*i*_ be the *i*-th (*i* ∈ 1, 2, *…, m*) feature vector. Each convolution operation in our model involves a filter, *w* ∈ ℝ ^*h×k*^, which is applied to a window of *h* features (*x*_*i*_, *x*_*i*+1_, *…, x*_*i*+*h−*1_) to produce a new refeature *c*_*i*_. And we can obtain a refeature map through convolution operation, *C* = [*c*_1_, *c*_2_, *…, c*_*m−h*+1_]. In our model, we also add several random feature embedding layers (i.e. 4) based on the original input integrated feature vectors, which are obtained from DeepMNE, to eliminate the effect of feature order. Therefore, the refeature map *C* ∈ ℝ ^*m−h*+1^ of convolution operation using single filter can be described as:

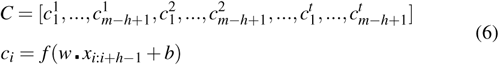

where *t* is the number of random feature embedding layers, *b* is a bias term and *f* is a non-linear function. After convolution operation, *c*_*i*_ is passed through a activation function (i.e. ReLu function *f* (*x*) = *max*(0, *x*)) that ignores negative outputs and propagates positive outputs from previous layer, because a higher *c*_*i*_ indicates that this captured refeatures of the feature combination very well. ReLU activations are used widely in deep learning model because of its computational efficiency and sparsity (Nair and Hinton, 2010), thus we also utilize in our CNN-based gene function prediction model. Besides, we also add batch normalization (Ioffe and Szegedy, 2015) layer in our model to accelerate training.

A max-pooling operation over the refeature map *C* ia utilized in our model, and we take the maximum value 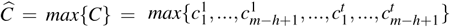 as the refeature corresponding to this particular filter, which can capture the most important refeature for each refeature map. Note that we use multiple filters in our model to obtain multiple refeatures in our model. Finally, these obtained refeatures are transferred to a fully connected *sigmoid* layer to obtain the final probability distribution over multi-labels.

We employed a constraint on *l*_2_-norms of the weight vectors on the penultimate layer (Hinton *et al.*, 2012). Besides, we also adopted dropout operations to prevent overfitting or co-adaptation of hidden units during forward-back propagation.

### 2.3 The DeepMNE-CNN algorithm

The core of DeepMNE algorithm is the semi-supervised autoencoder, and the goal of this model is to minimize the loss function Eq. (5). In this part, we will calculate the partial derivative of 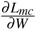 to validate the aforementioned model. Thus, the loss function of integrating constraints (Eq. 4) can be rephrased as follows:

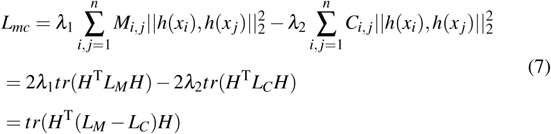

where *L*_*M*_ = *D*_*M*_ – *M, D*_*M*_ ∈ ℝ ^*n×n*^ is a diagonal matrix, 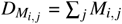. And *L*_*C*_ is similar as *L*_*M*_. *H* is the simplified representation of hidden layer. Thus, 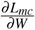 can be translated as:

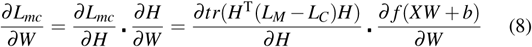

where *f* is activation function(i.e. sigmoid or ReLU), and we can obtain the partial derivatives of *L*_*MC*_. Thus, with a random initialization of the parameters, the proposed semi-supervised autoencoder can be optimized by using stochastic gradient descent (SGD).

The pseudocode for DeepMNE-CNN is given in Algorithm 1.

In the first phase of DeepMNE algorithm, we run random walk with restart algorithm to learn global structure of single biological network. Then, semi-autoencoder is trained to learn low-dimensional feature and extract prior constraints based on the network representation of hidden layer. Finally, CNN is employed for gene function prediction.

In the feature learning phase, DeepMNE algorithm uses an iterative model to train semi-supervised autoencoder with prior constraints. In each iteration, DeepMNE mainly contains three steps: merging constraints, training semi-autoencoder and extracting novel constraints. With the increasing of iterations, the model tends to converge and the constraints tends to be unchanged. Then, DeepMNE generates several low-dimensional feature representations of nodes. The DeepMNE algorithm is a scalable framework model, its training complexity is linear to the number of vertexes *N*. The part of extracting constraints need to calculate pair-wise PCC value which requires *O*(*N*^2^). Therefore, the training complexity of DeepMNE algorithm is *O*((*N*^2^ + *N*)*TK*), where *T* is the number of iteration and *K* is the number of multiple-networks.

#### Algorithm 1 The DeepMNE-CNN algorithm

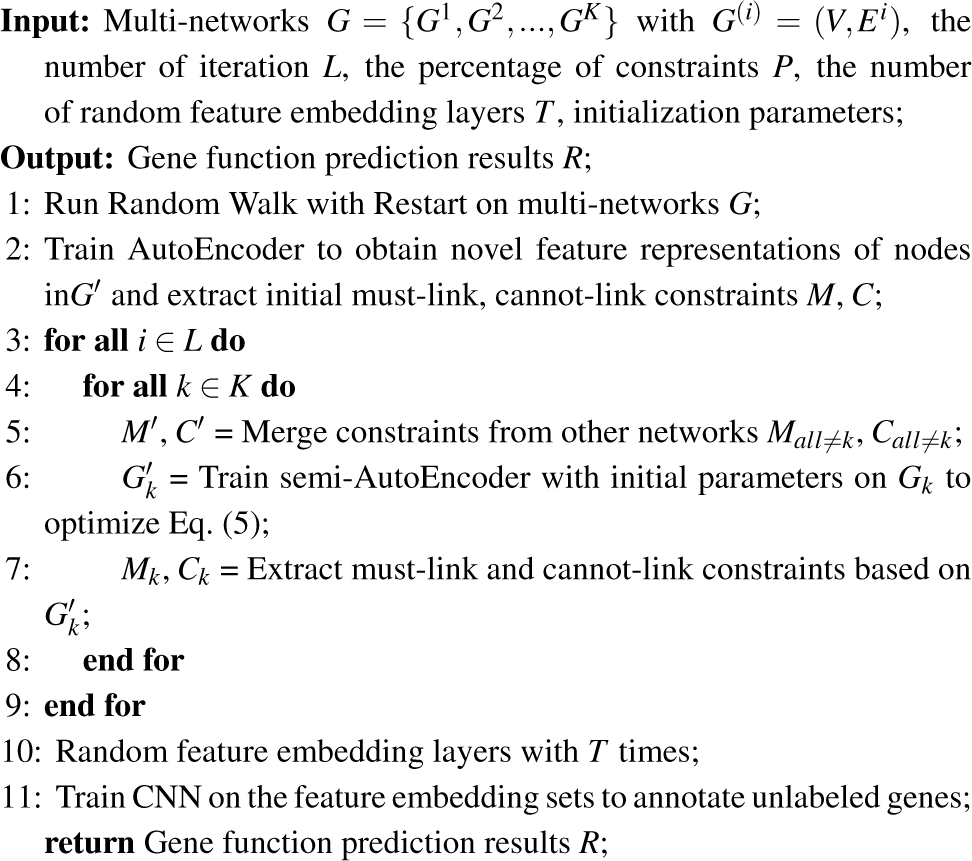

## 3 Results

### 3.1 Data preparation

To evaluate the performance of multi-networks integration framework (DeepMNE) and CNN-based gene function prediction, we implemented our method on Yeast and Human datasets respectively, which collected from the STRING database v9.1 (Franceschini *et al.*, 2013).

For yeast and human, the same dataset is also used in Mashup (Cho *et al.*, 2016). Yeast dataset mainly contains six networks with 6,400 gene nodes. The number of edges among different networks are between 1,1361 and 314,013. The edge weight in these networks varying from 0 to 1, which present the probability of edge presence. The functional annotations were downloaded from MIPS (Ruepp *et al.*, 2004). In detail, the functional categories in MIPS are grouped into three hierarchies, including Level-1, Level-2, and Level-3. In order to keep the independence among different categories, we removed categories whose Jaccard index score no less than 0.1 with other categories within the same hierarchy iteratively.

Human dataset also contains six networks with 18,362 genes, while the number of edges in these networks varying from 3,717 t0 1,544,348. The functional annotations were downloaded from the Gene Ontology database (M *et al.*, 2000). By grouping gene ontology terms, we can get three different functional categories, where each category contains 11-30, 31-100 and 101-300 genes respectively.

### 3.2 Parameters Setting

We compare our method, DeepMNE-CNN, with two state-of-the-art algorithms, Mashup (Cho *et al.*, 2016) and SNF (Wang *et al.*, 2014), to evaluate the performance on the task of annotating gene function. Function prediction can be treated as a multi-label classification task. Therefore, we adopt accuracy, micro-averaged F1, micro-averaged AUPRC and micro-averaged AUROC as the evaluation metrics. We randomly hold out 10% of the whole labeled genes as the validation set, 10% as the test set and utilized the remaining 80% as the train set to annotate unlabeled gene function. We repeatedly ran each method five times and adopt the average value as the final experimental results.

In the part of DeepMNE, parameters vary with different datasets. The dimension of each layer on multi-networks integration framework (DeepMNE) is listed in the supplementary document.

In the RWR part of our model, we use the same restart probability value as Mashup, which is 0.5. The final dimension of network representation are 500 and 800 respectively. The whole DeepMNE-based multi-networks integration algorithm is optimized by using stochastic gradient descent (Bottou, 1991). The batch size is 128 for yeast and 256 for human, the initial learning rate is 0.1 for yeast and 0.2 for human, and the epochs are 200 and 400 respectively. In the part of CNN-based gene function prediction, we add a dropout layer and batch normalization layer. We also employed l2-norms of the weight vectors. Parameters varies from different mini-datasets. For yeast level-1, the batch size is 64, learning rate is 0.001, the value of dropout is 0.5 and the epochs is 300. We utilized Adadelta optimizer to optimize the cnn model. For 31-100 category of human mf, the batch size is 64, learning rate is 0.01, the value of dropout is 0.5, the parameter of l2-norms is 0.005 and the epochs is 3000. We also utilized Adadelta algorithm to optimize the model.

For SNF, we generate an ensemble network and we run singular value decomposition to learn low-dimensional feature representation, and the dimension is same with our model.

### 3.3 Performance evaluation on Yeast datasets

We apply DeepMNE-CNN on three distinct functional categories of yeast dataset (level 1 with 17 categories, level 2 with 74 categories and level 3 with 154 categories) to validate its performance.

Comparing DeepMNE-CNN algorithm with two other approaches, we can observe consistent improvement on yeast dataset at all three annotation levels (see Figure 2). On the level-1 category of yeast dataset, DeepMNE-CNN labeled around 0.8378 of genes to their correct functional categories, in contrast to 0.8063 for Mashup and 0.6734 for SNF. Besides, the micro-F1 value of DeepMNE-CNN algorithm is 0.7096, which is sightly higher than other two methods, 0.6921 and 0.5957 respectively. The micro-average AUPRC and AUROC achieved by DeepMNE-CNN on level 1 of yeast are 0.7405 and 0.9100 respectively, which are significantly higher than the scores of Mashup and SNF. The detailed PRC and ROC figures of DeepMNE-CNN algorithm implemented on yeast can be found in the supplementary document.

**Fig. 2.**
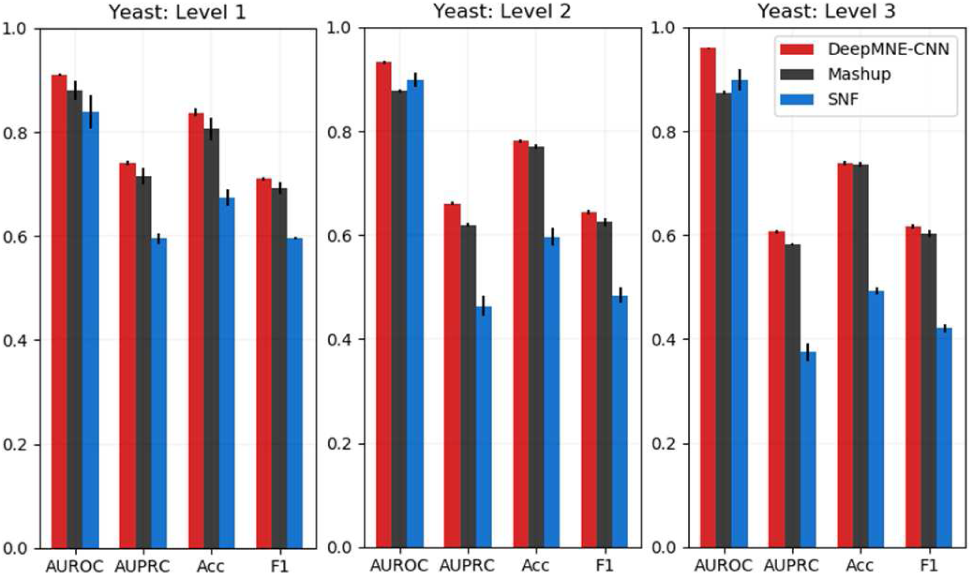
DeepMNE-CNN improves gene function prediction performance in yeast. We compared the function prediction performance of DeepMNE-CNN to other state-of-the-art approaches, Mashup and SNF in yeast. Figure represents the three different levels of functional categories respectively.

We also implement DeepMNE-CNN algorithm on each individual networks without integration and compared these AUPRC values with integrated network feature embedding on yeast-level-1 dataset (see Figure 3). We observed that our method significantly outperforms Mashup on single network except for cooccurrence and fusion network. Besides, we find that integrating multi-networks contributes to improving the performance of function prediction, which underscores the importance of integrating multiple types of data from various sources.

**Fig. 3.**
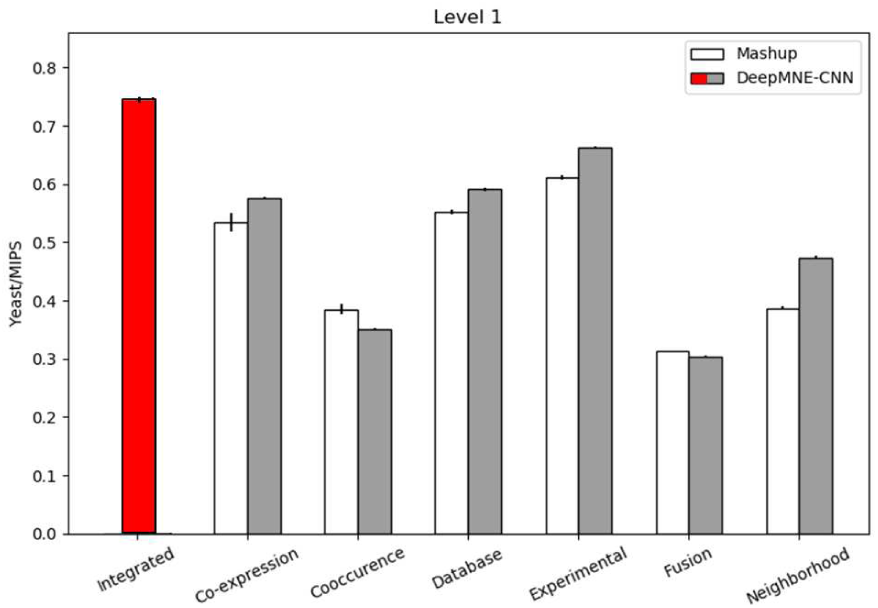
Integrating multi-networks performs better than individual networks in function prediction. The red shaded is the performance of DeepMNE-CNN implemented on integrated networks and the gray shaded is the AUPRC value of each individual network. The white one is the performance of Mashup.

### 3.4 Performance evaluation on Human datasets

For further evaluation, we also implement DeepMNE-CNN algorithm on human dataset to evaluate its performance. As described in the previous section, human covers three domains (BP, CC and MF) and three levels of functional categories. Thus, we can obtain nine distinct mini datasets and implement DeepMNE-CNN algorithm on these mini datasets to evaluate the performance.

Figure 4 shows the significant improvement of DeepMNE-CNN implemented on human Molecular Function and Cellular Component dataset. The accuracy of DeepMNE-CNN on human CC-101-300 is 0.5882, which is higher than Mashup and SNF (0.5626 and 0.4121 respectively). On human MF-101-300 mini dataset, DeepMNE-CNN still achieves the highest accuracy (0.5813) and significantly higher than the other two methods (Mashup, SNF are 0.5484 and 0.4137 respectively). The AUPRC values of DeepMNE-CNN implement on three categories of human MF are 0.5263, 0.3649 and 0.4007, which are all higher than two other methods (0.5150, 0.3636 and 0.3656 for Mashup, 0.3410, 0.1609 and 0.1748 for SNF). Besides, the AUROC values of DeepMNE-CNN (0.8280, 0.8488 and 0.8222 respectively) are all significantly higher than Mashup (0.8135, 0.8147, 0.8154) and SNF (0.7384, 0.7379, 0.7456). The detailed experimental results of DeepMNE-CNN on Biological Process (BP) dataset are listed in the supplementary document. ROC and P-R curves of nine mini datasets are also shown in the supplementary document.

**Fig. 4.**
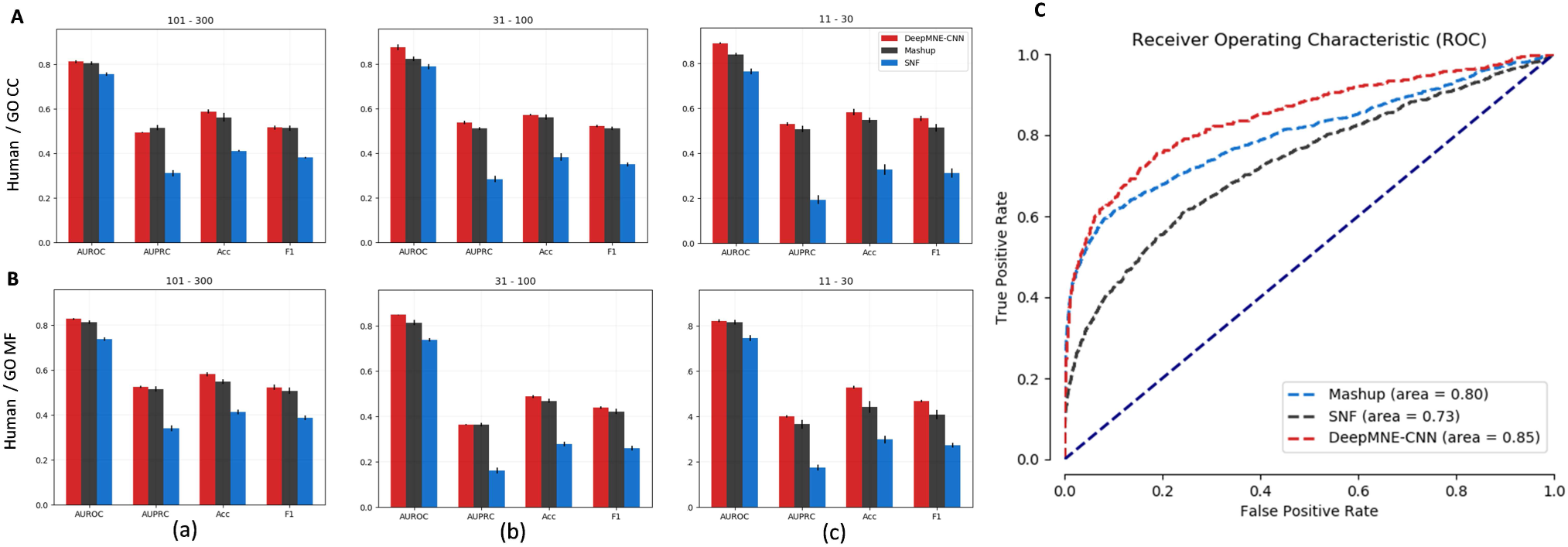
DeepMNE-CNN improves the performance of gene function prediction in Molecular Function and Cellular Component. We compared the function prediciton results of DeepMNE-CNN with other two state-of-the-art approaches, Mashup and SNF in human dataset. Figure A and B are evaluation results of MF and CC. Figure C displays the roc of 31-101 level of MF category on human.

### 3.5 Effects of DeepMNE-CNN components

The deep learning-based gene function prediction approach proposed on this paper mainly contains two parts, multi-networks integration algorithm (DeepMNE) and CNN-based annotation of gene function. In order to evaluate the performance of this two different parts, we implement a multi-networks integration framework without constraints sharing, called MultiAE-CNN, to evaluate the effects of proposed semi-AE. Furthermore, we implement SVM algorithm based on the integrated results of DeepMNE to predict gene function instead of CNN, termed as DeepMNE-SVM, which could evaluate the effectiveness of CNN on the task of annotating gene function. Besides, we also run Convolutional Neural Network (CNN) on multiple networks directly to predict gene function, which could validate the contribution of DeepMNE algorithm. Finally, we compare our method, DeepMNE-CNN, with these variational approaches to validate the effects of two distinct parts and the experimental results have been listed in Table 1.

**Table 1.**
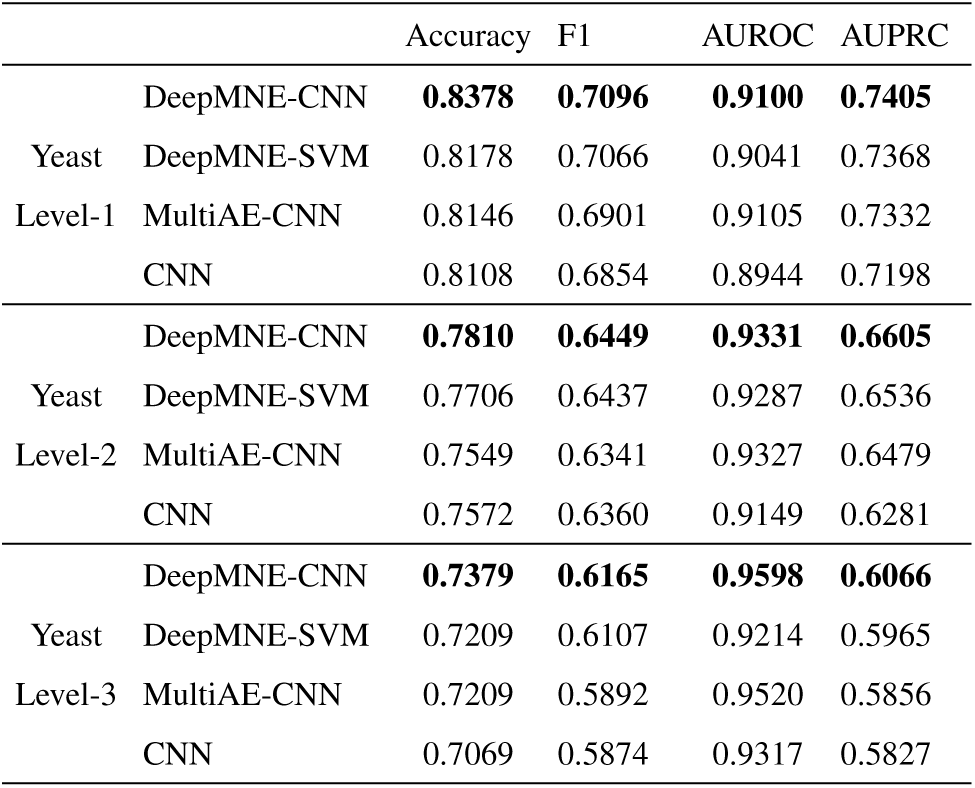
The Accuracy, micro-F1, AUROC and AUPRC of DeepMNE-CNN, DeepMNE-SVM, MutiAE-CNN, CNN on gene function prediction on yeast dataset.

The result shows that two parts of our proposed DeepMNE-CNN algorithm shows substantial superiority on the task of predicting gene function. Comparing with Mashup, which mainly contains network integration and SVM-based gene function prediction, DeepMNE-SVM indicates great performance in multi-networks integration (DeepMNE algorithm), where four evaluation indexes of DeepMNE-SVM are all significantly higher than Mashup. We also can find the superiority of DeepMNE algorithm by comparing with CNN. Besides, CNN-based gene function prediction also show great performance on the task of annotation of gene function by comparing DeepMNE-CNN with DeepMNE-SVM method. For instance, top prediction based on DeepMNE-SVM correctly labeled 0.8378 percentage of genes to true function categories on the level 1 of yeast dataset, where DeepMNE-SVM, MultiAE-CNN, CNN are 0.8178, 0.8146, and 0.8108 separately.

The restart probability of RWR and the dimension of embedding are critical parameters for the whole function prediction model. In order to evaluate the effects of these parameters to DeepMNE-CNN, we re-run our model with different number of restart probability and embedding dimension respectively. Figure 5. a shows that the AUPRC and AUROC are stable when varying embedding dimension from 100 to 900. Besides, the value of restart probability has a few effect on the performance of DeepMNE-CNN (see Figure 5. b). It is shown that DeepMNE-CNN is stable and robust, the effect of restart probability and embedding dimension on DeepMNE-CNN can be ignored.

**Fig. 5.**
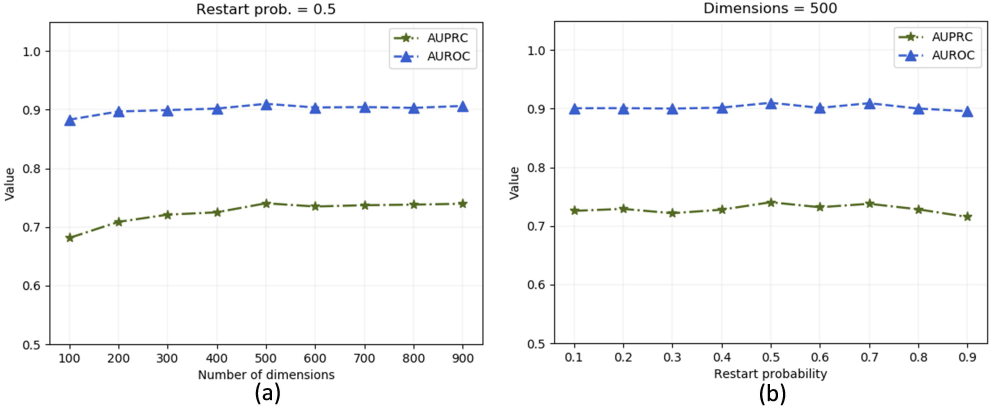
The AUPRC and AUROC score of DeepMNE with different restart probabilities and numbers of dimensions for function prediction on yeast dataset.

## 4 Conclusion

In this paper, we propose a novel multi-network integration algorithm, termed as DeepMNE, and apply it on gene function prediction using CNN model. We first extract constraints from various networks and use the semi-autoencoder to integrate different networks. Then, we utilize a convolutional neural network to predict gene function based on the feature vectors learned from DeepMNE framework. To demonstrate the performance of DeepMNE-CNN, we compare our method with two state-of-the-art measures. The evaluation on two real-world datasets demonstrates that DeepMNE-CNN performs better than other existing state-of-the-art approaches. Furthermore, we test the contribution of each component of DeepMNE-CNN. Overall, all evaluation tests show that DeepMNE-CNN implies great performance on multi-networks integration and function prediction.

